# Selection Of Theaflavin 2B As a Potential HSP90 Antagonist Semi-rigid Ligand Docking Analysis

**DOI:** 10.1101/2022.06.24.497557

**Authors:** Samuel Akinsehinwa, Oluwatobi Medayedupin

## Abstract

Heat shock protein 90 (HSP90) is an abundant molecular chaperone playing critical role as mediator of proper folding, maturation and stability of diverse client cellular proteins and has been reported to be overexpressed at a level of 2-10 folds relative to 1-2 folds of healthy cells. Geldanamycin and its derivatives (17AAG and 17 DMAG) has been developed to combat this but are known to be associated with primary side effects including plague, nausea, vomit, liver toxicity, hence the need to discover a relative safe and more potent inhibitor.

The aim of this study is to determine the inhibitory potential of the Theaflavin 2B, a furanocoumarin derivative, which has been documented to have anti-tumour activity against human prostate carcinoma DU145 cells via computational techniques. To this effect, theaflavin 2B was retrieved from PubChem database, and screened against HSP90 for its inhibitory effect, which resulted in relatively higher binding affinity of -9.2Kcal/mol relative to that of the standard (−6.4 Kcal/mol). Computational docking analysis were performed using PyRx, AutoDock Vina option based on scoring functions and the target was validated so as to ensure that the right target and appropriate docking protocol was followed for this analysis.

The docking studies of Theaflavin 2B with HSP90 showed that this ligand is a druggable molecule which docks well with HSP90 target. Therefore, theaflavin 2B molecule is a very good candidate for inhibiting HSP90 and presented it as good candidate for further evaluation as a potential therapeutic agent for cancer therapy.

---

The positive association between the recurring poor prognostic drug resistance and their varied genomic mutations widely emphasizes its clinical relevance in the various chemotherapeutic strategies for reducing cancer-associated morbidities and mortalities (66,67). This include the molecular chaperone HSP90.

The deregulation of HSP90 is an active driver of several oncogenic conditions including the highly untreatable KRAS-mutant lung adenocarcinoma (72,73).

We intend to find novel selective inhibitor (s) of HSP90, through molecular modelling, in this study that can effectively inhibit HSP90 with better potency and safety profile among the compounds reportedly identified in several studies (32,22,37,35), from various parts of *Moringa oleifera* – a scientifically proven anticancer plant in several tumour cell lines (36,27). This concept is validated by the fact that natural products are constituted of complex chemotypes with favourable safety profile and promising potential for discovery of novel HSP90 inhibitors (77).

## 2.0 Methods

### 2.1 Protein Preparation

The 3D coordinate files of crystal structure of an unliganded human (homo sapiens) HSP90 (PDB ID: 6X6O) was retrieved from the Research Collaboratory for Structural Bioinformatics (RCSB) protein data bank (http://www.rcsb.org), visualized using the Schrodinger molecular graphics program PyMol® and prepared using Schrodinger maestro protein preparation wizard. Briefly, the Ran-RanBP1 complex (chain A and B) were deleted while the remainder (chain C) was prepared under OPLS2005 force field at pH of 7.0±2.0. All steric clashes were corrected, hydrogen bond order fixed, while the missing side chains and loops were corrected using prime. Disulfide bonds were created, water molecules beyond 5.00Å were removed from the ionized (Het) groups, waters with less than 3 hydrogen bonds to non-waters were deleted while the bond orders were assigned. The charge cutoff for polarity was 0.25. This was further minimized by addition of polar hydrogens and merging of non-polar ones using the plugged-in version of open babel software of PyRx AutoDock Vina version 0.8 and Gasteiger charges were computed using AutoDock 4.226.

### 2.2 Ligand Preparation

The simplified input line entry system (SMILES) representations of identified *Moringa oleifera* compounds (32,41, 28,55) was obtained from PubChem database (https://pubchem.ncbi.nlm.nih.gov), while those with unavailable information were designed using ChemAxon Marvin Sketch. These were co-catenated and converted to MDL MOL structure description file format (SDF) using DataWarrior version 5.2.1. The physicochemical properties of the compounds were calculated and filtered based on the cut-off involving Lipinski’s rule, topological surface area (TPSA) ≤ 140Å, as well as the number of rotatable bonds ≤ 10. The compounds were also filtered further based on their putative tumorigenicity, mutagenicity, irritant properties, as well as druglikeness while the remaining compounds were converted to PDBQT file using PyRx tool to generate atomic coordinates through the optimization algorithm set at the universal force field parameters. The 2D SDF format of the standard compound (Geldanamycin) evaluated in this study was also obtained from PubChem and prepared with similar workflow.

### 2.3 Ligand Docking

We calculated the binding energies of the obtained compounds with the target binding site by single blind docking simulation using PyRx AutoDock Vina version 0.8 with further energy minimization through the forcefield-based scoring function (72). The PyRx AutoDock Vina, operating on the basis of assumably fixed three-dimensional target and flexible ligand structure (52), employs the Broyden-Fletcher-Goldfarb-Shanno method in its multithread gradient optimization approach of iterated local search global optimizer algorithm (44,7). The grid box resolution was centered at 12.0200 × -38.4570 × 337.6422 along the x, y and z axes respectively at the grid dimension of 25 Å x 25 Å x 25 Å defining the binding site and the comparative analysis of resulting interaction between the obtained *Moringa* compounds and the standard at the target binding site were obtained within the same docking grid box dimension.

### 2.4 Visualization of Docking Results

The ligand interactions at the target binding pocket and their three-dimension (3D) interaction with the target residues were visualized using Schrodinger Python Molecular graphics interface (PyMol®) version 2.2.0 software while the two-dimension (2D) ligand – residue interactions were obtained using BIOVIA Discovery Suite version 17.2.0.16349. The local minimum energy conformation binding mode of the compounds are regarded as the optimal binding pose at the target binding site in this study (56,46).

### 2.5 Validation of Docking Study

#### 2.5.1 Validation of Docking Protocol

The docking protocol of this study was validated by redocking of the same standard compound into the same target binding site and examining the relative binding pose of the ligands using the Python Molecular graphics interface (PyMol®) software.

#### 2.5.2 Validation of Docking Algorithm

The binding energy calculation algorithm applied in this study was validated by comparative analysis of docking scores of the hits and the standard compound (KPT-185) in other different docking algorithms including Molegro virtual docker (PLANTS scoring function), LeDock, and iGEMDock.

## Results and Discussion

Heat Shock Protein 90 has stabilizing effect on the conformations of mutant proteins during transformation, favours the emergence of polymorphisms and mutations that support the evolution of resistant clones; this is a rationale for targeting HSP90-dependent pathways in cancers and inflammatory diseases (Servin *et al*, 2015).

Theaflavin 2B, the lead compound has a binding energy of -9.2kcal/mol, while the standard compound has binding energy of -6.5 kcal/mol (Table 1). The highest binding energy (−9.2kcal/mol) attributed to Theaflavin 2B (PUBCHEM ID: 5441067) in this regard is believed to be as a result of its higher chemical interactions at the receptor active site; thirteen interactions in total: Twelve (12) **(Table 2; figure 3a)**, Hydrophobic interaction involving L-107, F-138, W-162, F-22, F-170, M-98, V-150, L-107 residues and one Hydrogen bond involving Y-139 residue, while, that of the standard ligand (geldanamycin) presents with the following chemical interactions at the binding pocket ***(Table 3, figure 3b)***. Two (2) hydrogen bonds involving T-184 and G-135residues ; Eight (8) hydrophobic interactions involving A-55, (2) M-98, L-107, F-138, L-48, V-186, A-55.; Two (2) other interaction involving M-98 residue.

**Table 1:**
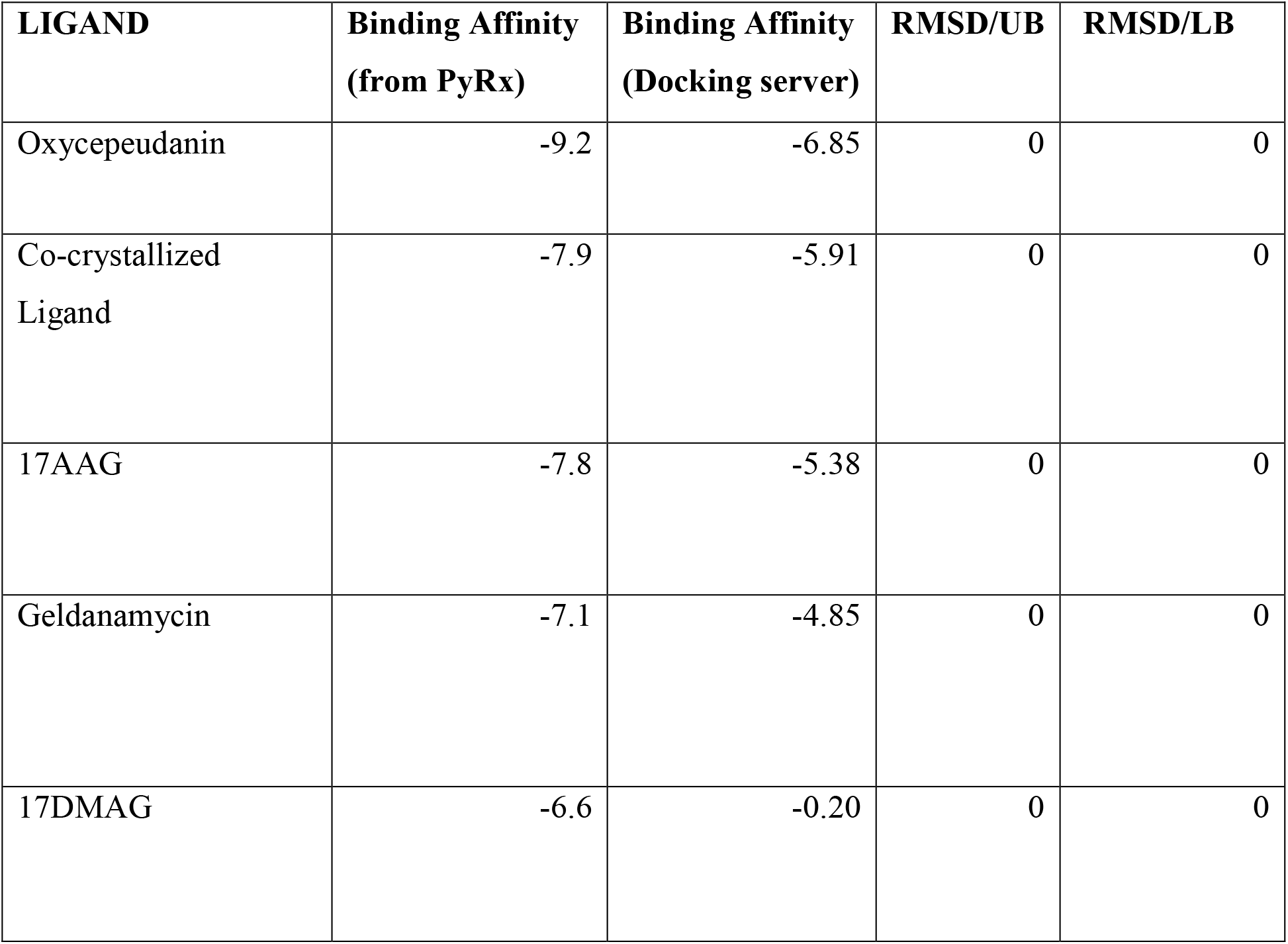
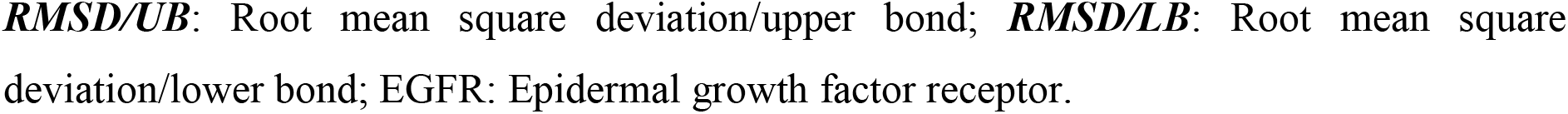
Energy and RMSD values obtained during docking analysis.

**Table 2:** Interaction table showing the chemical interactions of the co-crystallized within the binding pocket.

| NAME | CATEGORY | TYPES |
| --- | --- | --- |
| A:T184:HG1 - N:KFY:N | Hydrogen Bond | Conventional Hydrogen Bond |
| N: KFY:C - A:G135:O | Hydrogen Bond | Carbon Hydrogen Bond |
| A:A55:CB - N: KFY | Hydrophobic | Pi-Sigma |
| A:M98:CE - N: KFY | Hydrophobic | Pi-Sigma |
| A:M98:CE - N: KFY | Hydrophobic | Pi-Sigma |
| A:L107:CD1 - N: KFY | Hydrophobic | Pi-Sigma |
| A:M98:SD - N: KFY | Other | Pi-Sulfur |
| A:M98:SD - N: KFY | Other | Pi-Sulfur |
| A:F138 - N: KFY | Hydrophobic | Pi-Pi Stacked |
| N: KFY:C - A:L48 | Hydrophobic | Alkyl |
| N: KFY:C - A:V186 | Hydrophobic | Alkyl |
| N: KFY - A:A55 | Hydrophobic | Pi-Alkyl |

THR-T, GLY-G, ALA-A, MET-M, LEU-L, PHE-F, VAL-V.

**Table 3:** Interaction Table showing the various chemical interactions of Oxycepeudanin within the binding pocket.

| NAME | CATEGORY | TYPES |
| --- | --- | --- |
| A:Y139:HH -<br>N:OXYPEUCEDANIN:O | Hydrogen Bond | Conventional Hydrogen Bond |
| A:L107:CD1 - N:<br>OXYPEUCEDANIN | Hydrophobic | Pi-Sigma |
| A:F138 - N: OXYPEUCEDANIN | Hydrophobic | Pi-Pi Stacked |
| A:F138 - N: OXYPEUCEDANIN | Hydrophobic | Pi-Pi Stacked |
| N: OXYPEUCEDANIN - A:F138 | Hydrophobic | Pi-Pi Stacked |
| A:W162 - N: OXYPEUCEDANIN | Hydrophobic | Pi-Pi T-shaped |
| N: OXYPEUCEDANIN - A:W162 | Hydrophobic | Pi-Pi T-shaped |
| A:F22 - N: OXYPEUCEDANIN:C | Hydrophobic | Pi-Alkyl |
| A:F22 - N: OXYPEUCEDANIN:C | Hydrophobic | Pi-Alkyl |
| A:F170 - N:<br>OXYPEUCEDANIN:C | Hydrophobic | Pi-Alkyl |
| N: OXYPEUCEDANIN - A:M98 | Hydrophobic | Pi-Alkyl |
| N:OXYPEUCEDANIN - A:V150 | Hydrophobic | Pi-Alkyl |
| N:OXYPEUCEDANIN - A:L107 | Hydrophobic | Pi-Alkyl |

***TYP-Y, LEU-L, PHE-F, TYP-W, MET-M, VAL-V.***

**Figure 2:**
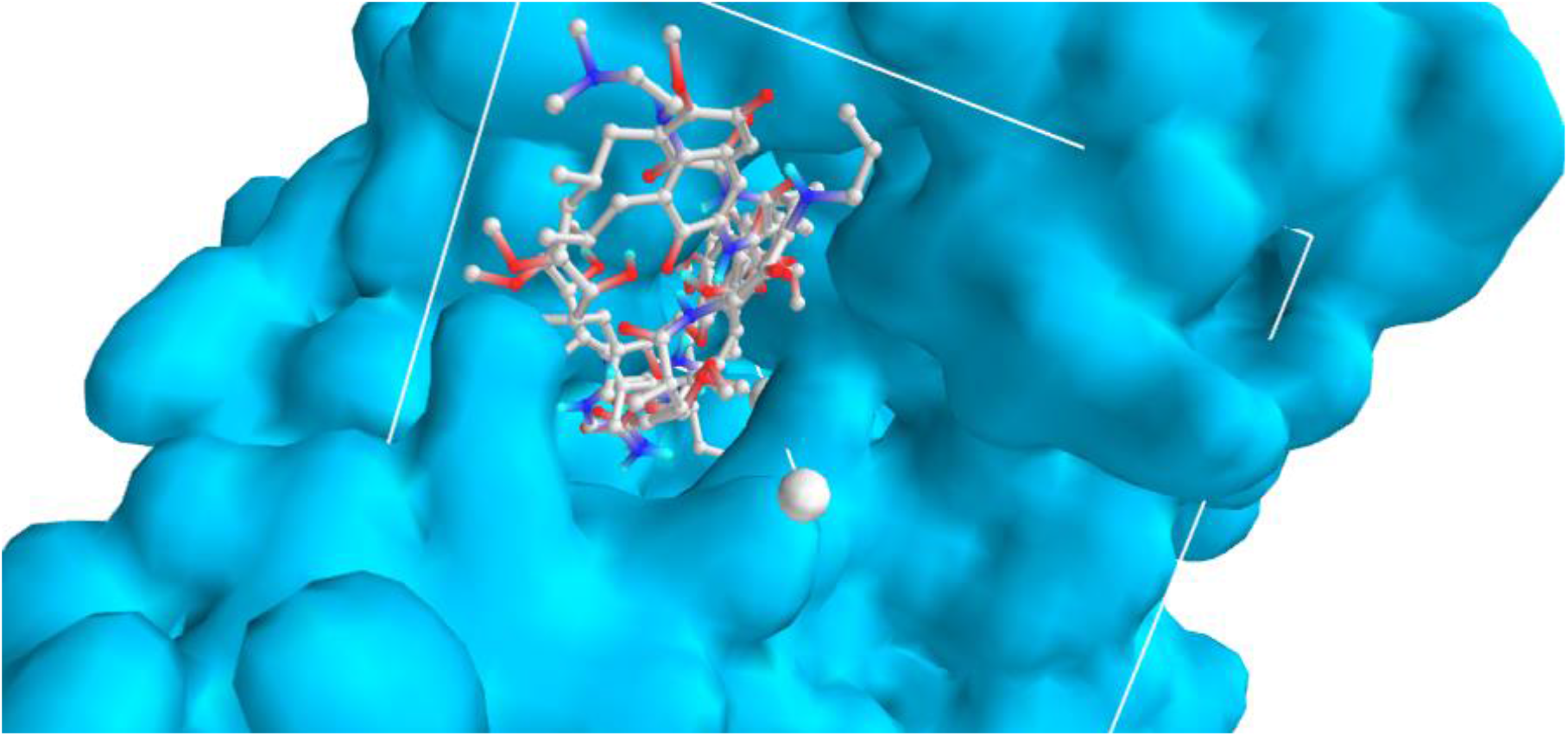
Grid box within which the ligand binds 0.3021 × 14.7950 × 216.4475 along the X, Y, Z-axis

**Figure 3:**
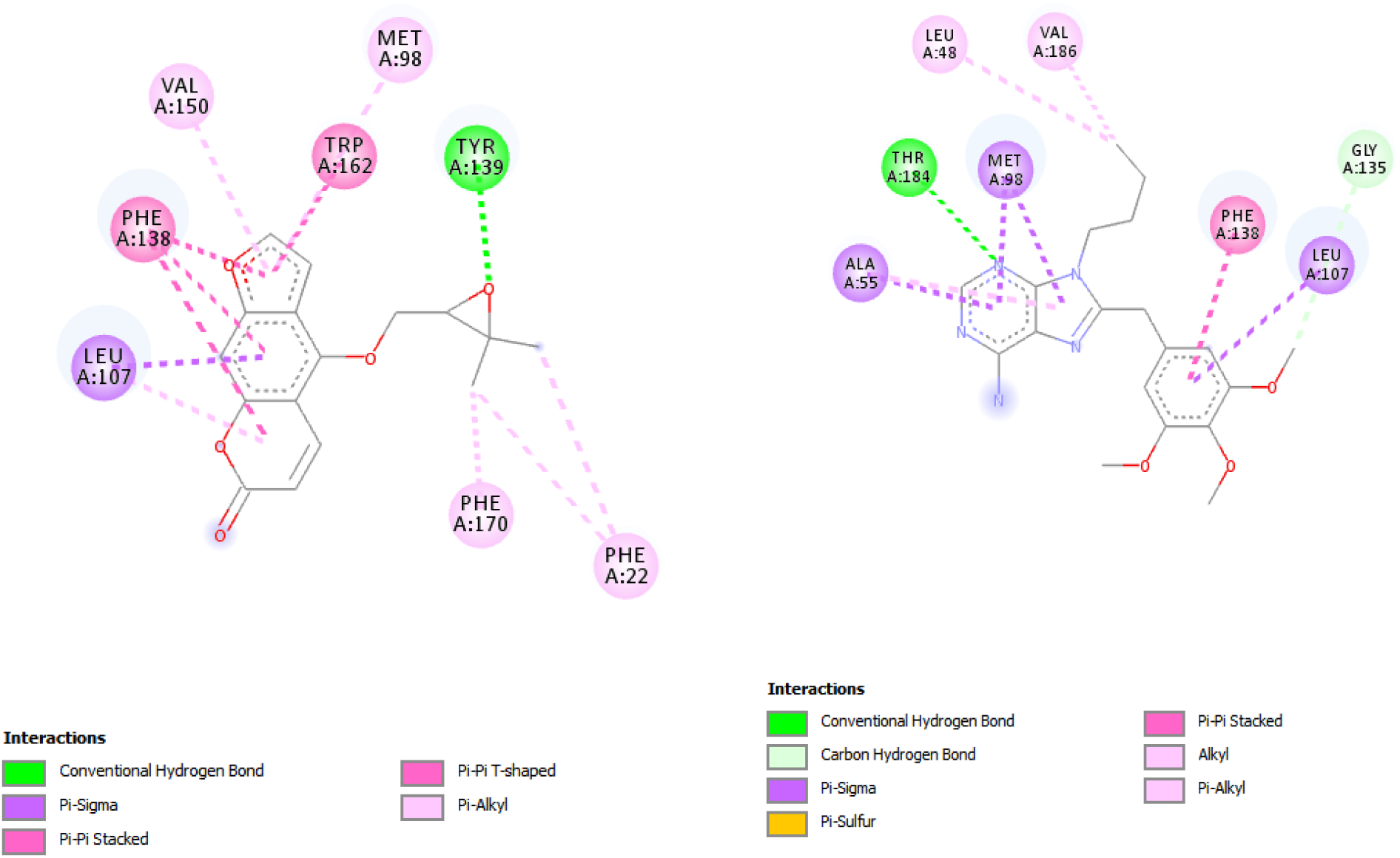
2D interactions of ligands within the binding pocket (a)Oxypeucedanin (b)KYP.

**Figure 4:**
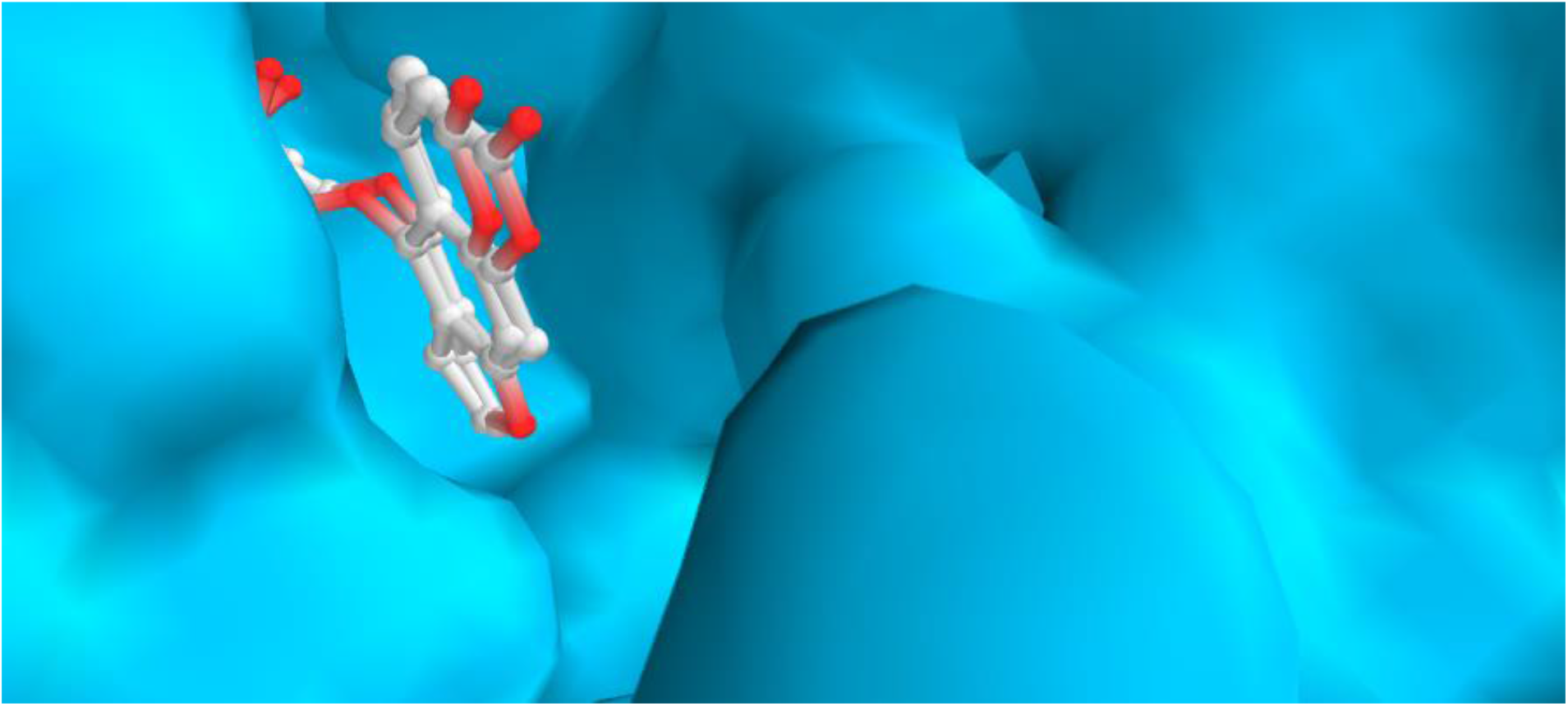
Validation of docking: Comparability of the re-docked binding mode and the lead compound-Oxypeucedanin pose with the accompany residues of HSP90 binding pocket. A snapshot from PyRx.

## Conclusion

The docking studies and ADMET evaluation of Theaflavin 2B with HSP90 showed that this ligand is a druggable molecule which docks well with HSP90 target. Therefore, Theaflavin 2B molecule plays an important role in inhibiting HSP90 and thus should be further evaluated as a potential therapeutic agent for cancer therapy.

## Notes

### Competing Interest Statement

The authors have declared no competing interest.

